# Spatially distinct phytohormone responses of individual *Arabidopsis thaliana* root cells to infection and colonization by *Fusarium oxysporum*

**DOI:** 10.1101/2022.12.20.521292

**Authors:** Jacob Calabria, Liu Wang, Madlen I. Rast-Somssich, Hsiang-Wen Chen, Michelle Watt, Staffan Persson, Alexander Idnurm, Marc Somssich

**Affiliations:** Plant-Fusarium Interactions Research Team, School of BioSciences, University of Melbourne, Parkville, Australia; Crop Root Physiology Lab, School of BioSciences, University of Melbourne, Parkville, Australia; Plant Cell Biology Lab, School of BioSciences, University of Melbourne, Parkville, Australia; Copenhagen Plant Science Center, University of Copenhagen, Frederiksberg C, Denmark; Mycology Laboratory, School of BioSciences, University of Melbourne, Parkville, Australia

**Keywords:** *Arabidopsis thaliana*, *Fusarium oxysporum*, *Fo*5176, salicylic acid, jasmonic acid, ethylene, plant immunity, spatial immunity, plant-microbe interactions

## Abstract

Jasmonic acid (JA), ethylene (ET) and salicylic acid (SA) are the three major phytohormones coordinating a plant’s defense response to pathogenic attack. While JA and ET are assumed to primarily control the defense against necrotrophic pathogens, SA-induced defense responses target mainly biotrophic microbes, and can include drastic measures such as programmed cell death as part of the plant’s hypersensitive response (HR). *Fusarium oxysporum* is a hemibiotrophic fungal pathogen of several plant species, including many important food crops, and the model plant species *Arabidopsis thaliana*. Colonization of the plant’s root vascular tissue by the fungus eventually results in wilting and plant death. A general role for JA, ET and SA in combating infection and colonization of the plant by *F. oxysporum* has been demonstrated, but their distinct roles and modes of action have so far not been described. Here, using high resolution microscopy with fluorescent marker lines of *A. thaliana* roots infected with *F. oxysporum* we show that SA acts spatially separate from JA, in a distinct set of root cells immediately neighboring the fungal colonization site. There, SA induces HR to stop the spread of colonization. JA acts in a different, but also clearly defined set of cells, slightly removed from the colonization site, where it initiates a defense response to actively resist the invader. ET is activated in a stretch of cells that covers both, the cells with activated SA and JA signaling, and may be involved in creating these two distinct zones. These results show how the three phytohormones act together, but spatially and functionally separate from each other, to fight this hemibiotrophic pathogen. Such a high-resolution analysis to resolve the plant’s immune response to pathogenic infection on an individual cell level and in intact tissue has so far been lacking.

**Graphical Abstract:**

- Colonization of the *A*. *thaliana* root tip by *F*. *oxysporum* strain *Fo*5176 leads to immediate cell death of the colonized and surrounding tissue.
- As the colonization front progresses through the vasculature, the cell death front moves along with it through not only the vasculature, but also the surrounding tissues.
- WRKY70 positively regulates salicylic acid (SA) biosynthesis in cells immediately adjacent to the colonized tissue, inducing a hypersensitive response (HR), thereby killing off the cells deemed lost to the intruder, establishing the cell death front.
- Slightly further removed from the HR zone, WRKY11 induces jasmonate (JA) biosynthesis in cells of the vasculature to launch a defense response aimed at actively repelling the fungus.

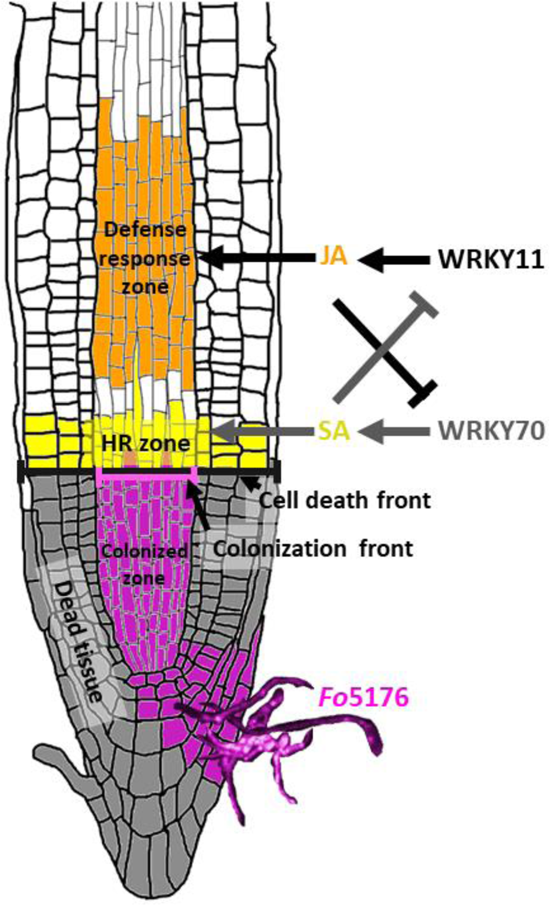

## Introduction

Phytohormones play an important role in coordinating a plant’s defense response to attack by pathogenic microbes (Hou and Tsuda, 2022). A plant senses and recognizes a potential attacker via their microbe-associated molecular patterns (MAMPs); highly conserved structures on the outside of the microbe, such as flagellin in the bacterial flagellum, or chitin in the fungal cell wall (Wang *et al*., 2022b). Since the root is continuously surrounded by microbes carrying MAMPs in the soil, recognition of a MAMP alone will not trigger the plant immune system. Instead, it is the co-incidence of MAMPs with cell damage caused by a pathogenic microbe that will launch a full immune response (Zhou *et al*., 2020). MAMPs and damage-associated molecular patterns (DAMPs) are sensed by the plant outside the cell via plasma membrane localized PATTERN RECOGNITION RECEPTORS (PRRs), which then transduce the information about the attacker into the cell via an intracellular kinase domain (Lee *et al*., 2021). Inside the cell, several defense pathways are then activated, including phytohormone-dependent signaling pathways (Lee *et al*., 2021). Jasmonic acid (JA), ethylene (ET) and salicylic acid (SA) are the three main phytohormones involved in coordinating the plant’s defense against pathogenic invaders (Hou and Tsuda, 2022). Among them, JA and ET mostly act in concert, while JA and SA appear to act in mutual antagonism. It is generally assumed that JA/ET control the plant’s defense against necrotrophic pathogens, while SA is the regulator of pathways combating biotrophic pathogens (Hou and Tsuda, 2022). SA-mediated immunity can involve drastic measures such as the hypersensitive response (HR), a form of programmed cell death aimed at containing an infection by killing off the tissue surrounding the infection site in an attempt to prevent further spread of the microbe to other healthy cells and tissues (Pitsili *et al*., 2020). This form of defense is not activated against necrotrophic pathogens, as these feed on dead plant material and would therefore benefit from such a cell death response (McCombe *et al*., 2022). Thus, the JA/ET-mediated response to necrotrophs instead focusses on active mechanisms that can prevent spread of the pathogen, for example, releasing reactive oxygen species (ROS) into the apoplast that can harm and kill the microbe, producing antimicrobial compounds, or changing the apoplastic pH (Masachis *et al*., 2016; McCombe *et al*., 2022).

*Fusarium oxysporum*, a soil-inhabiting ascomycete fungus, is a destructive pathogen to many plant species, including several major crop plants, such as the tomato (*Solanum lycopersicum*), cotton (*Gossypium barbadense*), and banana (*Musa acuminata*) (Takken and Rep, 2010; Gordon, 2017; Dita *et al*., 2018). *F. oxysporum* reproduces asexually and the resulting clonal lineages all represent individual formae speciales (ff. spp.), that together form the *F. oxysporum* species complex (Summerell, 2019). The different ff. spp. mostly share an 11 chromosome core genome and it is their lineage-specific accessory chromosomes that account for the host specificity of each f. sp., and whether a f. sp. is pathogenic, non-compatible, or beneficial to a certain plant (Ma *et al*., 2010; Redkar *et al*., 2022). *F. oxysporum* hyphae grow in the soil and are chemotactically guided toward the root of their host plant (Wang *et al*., 2022a). Upon reaching the plant, they attach and grow along the root until they enter via natural openings, such as lateral root emergence sites or wounds (Wang *et al*., 2022a). Inside the root, the hyphae grow into the vasculature, infecting and colonizing xylem vessel cells (van der Does *et al*., 2008). Once inside the vasculature, the fungus drains the plant of water and nutrients, and can spread to all parts of the plant, which begins to wilt and eventually dies (Gordon, 2017). Based on these observations, *F. oxysporum* is considered a hemibiotrophic fungus, that initially lives biotrophically within the host’s vasculature before eventually turning necrotrophic and killing the plant (Gordon, 2017). Research into the plant’s defense against *F. oxysporum* has originally focused on the crop plants threatened by Fusarium wilt (e.g. Panama disease of bananas) and the *F. oxysporum* ff. spp. targeting them. However, more recently, the *Arabidopsis thaliana* – *F. oxysporum* f. sp. *conglutinans* strain 5176 (*Fo*5176) pathosystem has also been established to study this plant-fungus interaction with all the modern molecular, cell biological, genetic and genomic tools available for this plant model system (Wang *et al*., 2022a).

Early work on the *A. thaliana* – *Fo*5176 pathosystem has focused on creating several large-scale transcriptomic datasets, as well as comprehensive infection studies of *A. thaliana* mutants, to investigate the role of JA and SA in combating infection by *Fo*5176 (Wang *et al*., 2022a). These studies have established a role for the different hormones in the plant’s defense against *Fo*5176, but were also often inconclusive as to their exact roles (Wang *et al*., 2022a). Thus, while it is assumed that both SA and JA are involved in the plant’s defense response against this hemibiotrophic fungus, a detailed understanding of their action is lacking (Wang *et al*., 2022a).

A possible explanation for the failure of large-scale transcriptomic approaches to identify novel and specific pathways involved in the defense against *Fo*5176, as well as the apparent inconclusiveness in relation to known pathways, such as the phytohormones, may lie in the tissue and timing used in the experiments. In contrast to other plants, such as tomato, where *F*. *oxysporum* can infect the plant via several openings simultaneously, the fungus generally appears to infect *A. thaliana* solely via the root tip, thereby limiting the number of infection sites and thus infected tissue available to be subjected to transcriptomic analyses (Czymmek *et al*., 2007; Wang *et al*., 2022a; Martínez-Soto *et al*., 2022). Thus, by using either whole seedlings or the whole root system to extract RNA, the transcriptional changes in these few infected cells may be diluted too much by the other uninfected cells. Furthermore, as the plant also senses the fungal hyphae in its rhizosphere via their MAMPs, a majority of root cells will launch a general MAMP-response, which further dilutes the response of the cells undergoing infection.

For our work, we have set up a microscopy-based live-imaging approach monitoring transcriptional activity of different pathways *in planta*, over time and on an individual cell level. We have previously published our pGG-PIP GreenGate entry vector collection with regulatory sequences for plant immunity genes, including reporters for the different phytohormone pathways (Calabria *et al*., 2022). Using this approach, we detected local responses of genes involved in SA, JA and ET biosynthesis, signaling and regulation, targeted to specific tissues and to a very limited number of root cells upon infection with *Fo*5176. Interestingly, we found that SA acts spatially separate from JA in a distinct set of root cells in direct contact with the fungus, where it appears to induce HR. JA in turn also acts in a clearly defined set of cells which are slightly removed from the colonization site and SA-induced HR zone, where it appears to induce the plant’s active defense against the pathogen.

## Results

### *Fo*5176 colonization of the root leads to cell death of the infected tissue

To monitor the infection and colonization of *A. thaliana* roots by *Fo*5176, we cultivated 10-day old seedlings in square petri dishes, added fungal spores and then imaged hyphal growth, infection and colonization over the next 14 days. We used a transgenic *Fo*5176 line expressing a cytoplasmic tdTomato fluorophore to visualize the fungus (Calabria *et al*., 2022). One of the first things noticeable when imaging infected plants was that the tissue surrounding the colonization site appeared to undergo immediate cell death. When cells die, the plasma membrane and vacuole collapse, leaving behind the cell wall material. As this material scatters light, the dead tissue appears as bright white in the image (Fig. 1A). In fact, when the vasculature of a root tip was colonized by *Fo*5176, the entire tip underwent cell death (Fig. 1A, B). By imaging a plant line expressing a fluorescent marker for the plant plasma membrane (PM) (mNeonGreen(mNG)-LTI6b), this became even more apparent, as the fluorescence signal from the PM outlined all cells of the plant but disappeared completely from all cells around the colonized tissue (Fig. 1C, D). As fungal colonization progressed through the vasculature as a ‘colonization front’, a ‘cell death front’ was also observed that moved in line with it. In contrast though, the cell death front spanned the entire circumference of the root, including the outer tissues, epidermis, cortex, and endodermis, which are not colonized by the fungus. Because cell death was not limited to the colonized tissue, we reasoned that cell death may be the result of a hypersensitive response (HR) initiated by the plant, rather than caused directly by *Fo*5176.

**Figure 1:**
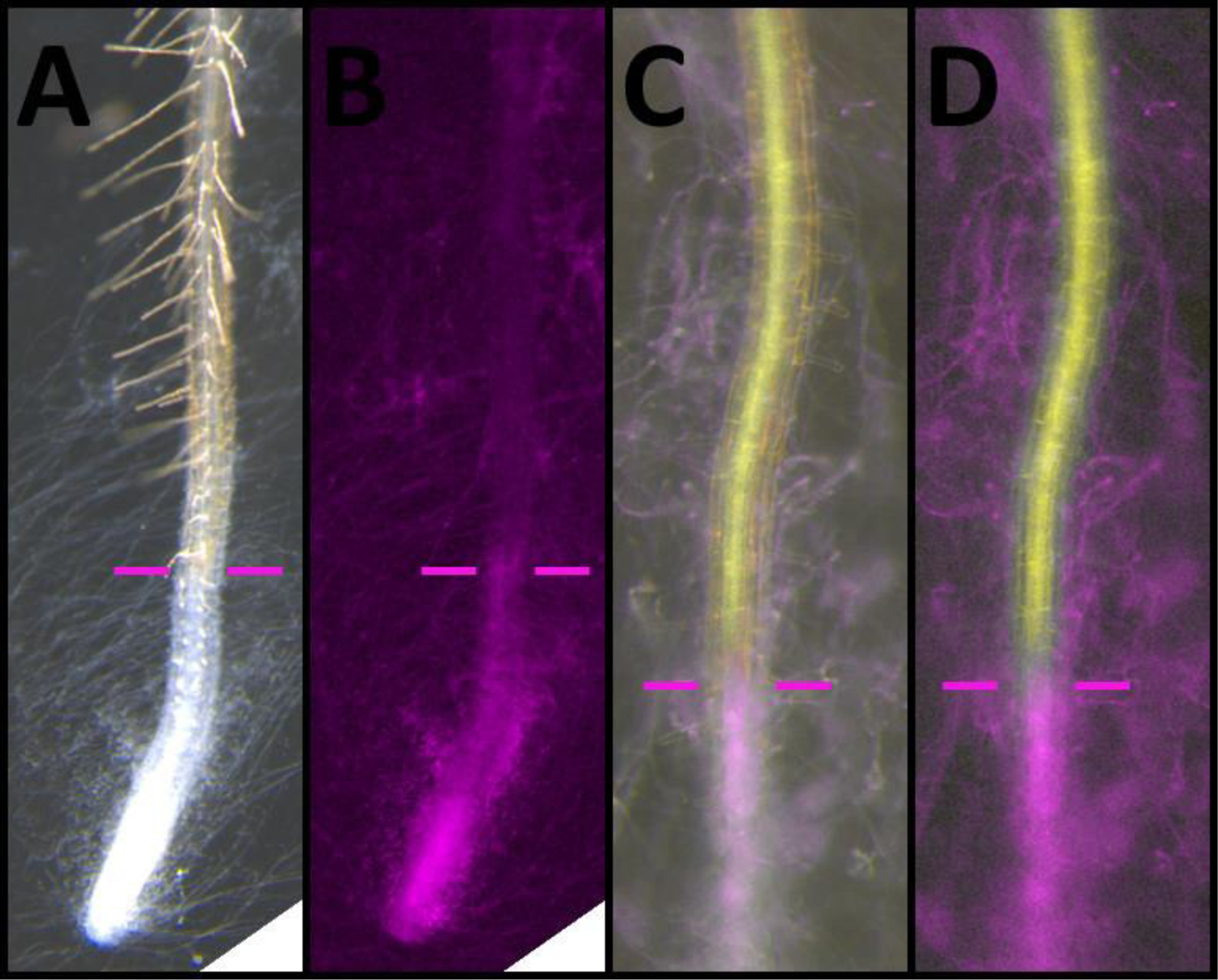
Colonization of *A. thaliana* roots by *Fo*5176 causes cell death. **A, B**: Bright field (A) and fluorescence image (B) of a colonized root tip. (A) Dead cells appear bright white. (B) Fluorescent fungal hyphae (magenta) can be seen in the tip and the vasculature. The colonization front (magenta bars to the left and right of the root) aligns with the cell death front. **C, D**: Overlay of bright field and fluorescence image (C) and fluorescence image only (D) of a root expressing the PM-marker mNG-LTI6b (yellow) and colonized by *Fo*5176 (magenta). Fluorescent hyphae (magenta) can be seen in the tip and the colonized vasculature. The fluorescence coming from the PM-marker (yellow) is lost at the colonization front (magenta bars).

### Salicylic acid signaling is activated in the hypersensitive response zone

As HR is often activated via an SA- and PATHOGENESIS-RELATED PROTEIN 1 (PR1)-dependent pathway, we next investigated if SA biosynthesis and signaling via the antimicrobial protein PR1 are indeed activated in the cells bordering the colonized tissue. We utilized reporters for the key SA biosynthesis enzyme ENHANCED DISEASE SUSCEPTIBILITY 16 (EDS16; also named SALICYLIC ACID DEFICIENT 2 (SID2)) and PR1. In an uninfected control, *EDS16* was expressed in the meristematic and elongation zones (MZ/EZ) of the root tip, as well as in older differentiated cells, with no expression detectable in young differentiating cells (Fig. S1A, B). In the tip, expression also appeared to be stronger in the outer layers, epidermis and cortex, compared to the vasculature (Fig. S1A, B). *PR1* expression was only weakly detectable in the mature differentiation zone (DZ; Fig. S1C, D). However, following infection and colonization by *Fo*5176, *EDS16* expression was upregulated in the root tip, and spatially targeted to a group of vascular cells immediately at the colonization front of the fungus (Fig. 2A, B). *PR1* expression, which was not detectable in the tip of the uninfected control, was also upregulated in these same cells, the set of cells likely to subsequently initiate HR (Fig. 2C, D).

**Figure 2:**
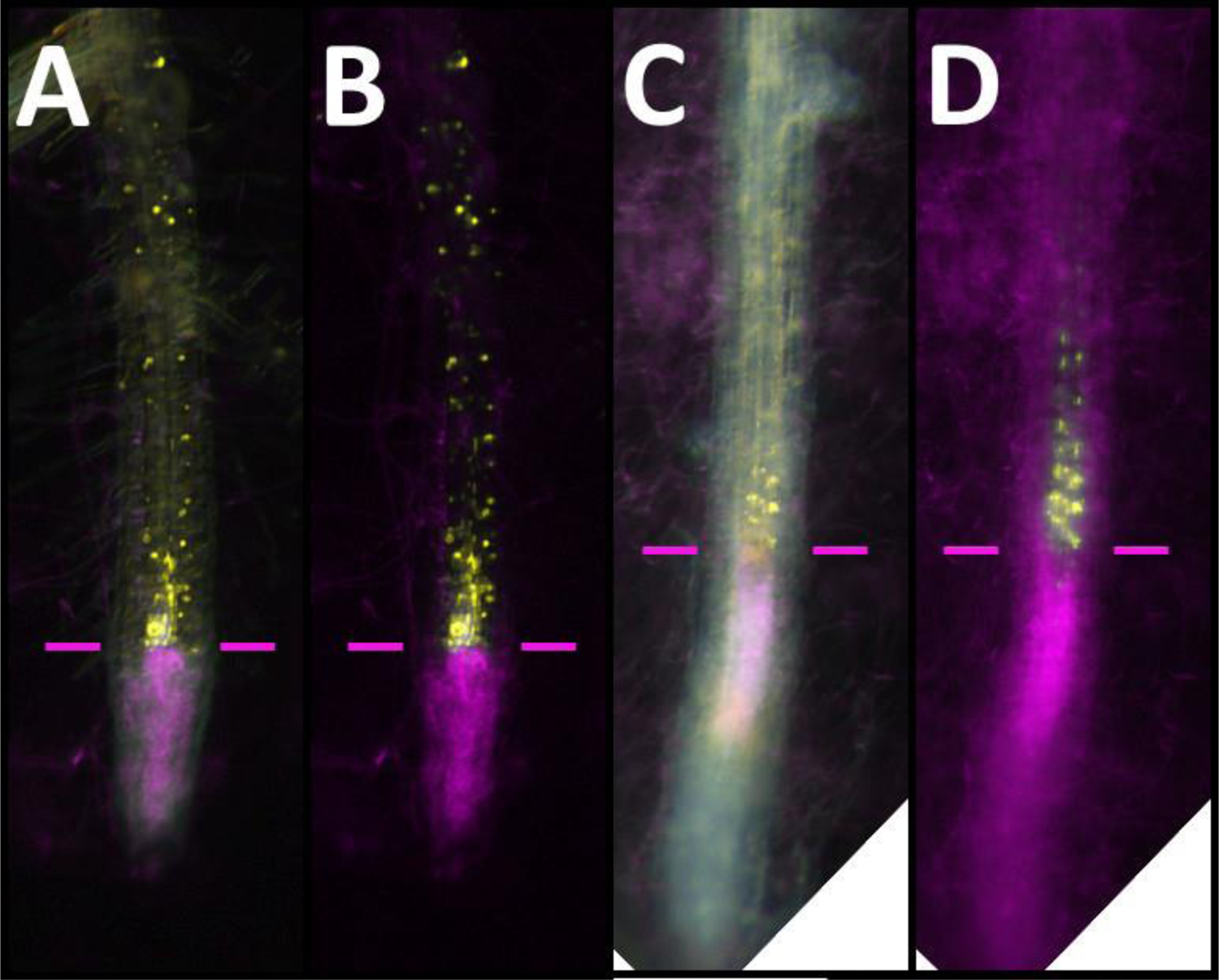
Expression of *EDS16* and *PR1* following colonization of the root by *Fo*5176. **A, B**: Overlay of bright field and fluorescence (A), and fluorescence-only (B) images of a root tip expressing the *EDS16* reporter (yellow) in response to colonization by *Fo*5176 (magenta). **C, D**: Overlay of bright field and fluorescence (C), and fluorescence-only (D) images of a root tip expressing the *PR1* reporter (yellow) in response to colonization by *Fo*5176 (magenta). The colonization front is marked by magenta bars.

### Jasmonic acid and ethylene act in a distinct group of cells close to the colonized tissue

We then investigated the responses of our JA and ET reporters. In the case of JA, we used regulatory sequences for *ALLENE OXIDE SYNTHASE* (*AOS*), a key enzyme in the JA biosynthesis pathway, *PLANT DEFENSIN 1.2* (*PDF1.2*), a defense gene activated as the result of active JA-signaling, and *ETHYLENE RESPONSE FACTOR 1* (*ERF1*), an activator of JA- and ET-responsive defense gene expression, as well as an integrator of the JA- and ET-signaling pathways (Wang *et al*., 2022a). To monitor ET-biosynthesis, we used our reporter for 1-AMINOCYCLOPROPANE-1-CARBOXYLIC ACID SYNTHASE 2 (ACS2), a key enzyme in the ET biosynthesis pathway. We found that *AOS* expression was mostly absent in the uninfected control roots, with very weak expression limited to sites of natural stress occurrence, such as lateral root emergence sites (Fig. S2A, B). Similarly, *PDF1.2* was not expressed in the root; however, it was strongly expressed in above ground tissue. *ERF1* was expressed in all differentiated tissues, but was absent from the root tip, including the MZ and EZ (Fig. S2C-F). This contrasting expression pattern between *AOS* and *ERF1* most likely represents activation via the ET-, rather than the JA-pathway, since ET also acts in several developmental pathways, while JA-signaling is mostly restricted to stress conditions. Indeed, the marker for ET-biosynthesis, *ACS2*, was expressed throughout the root system in the uninfected control, with the main root showing stronger expression than the lateral roots (Fig. S2G-J). Furthermore, expression was absent from the epidermis in differentiated tissues, and the expression was generally very weak between the EZ and early DZ. Expression in the MZ was moderate and included the epidermal cells (Fig. S2G-J).

Following infection and colonization of the plant by *Fo*5176, *AOS* expression was robustly induced in a distinct group of cells roughly four to seven cells away from the colonized tissue (Fig. 3A, B). The expression then slowly tapered out along the root toward the base. Interestingly, expression was limited to cells of the vasculature, with no upregulation in the epidermis, cortex, or endodermis. Thus, *AOS* expression and JA signaling are precisely targeted to the tissue that is colonized by the fungus. Interestingly, *PDF1.2* was not responsive in the group of cells next to the colonization site, despite this gene being routinely used as a general readout for JA-signaling activity in PCR or transcriptomic assays (Fig. 3C, D). However, at later stages of infection, when the fungus had progressed far enough along the vasculature to reach the DZ, either through the main root or along a lateral root, *PDF1.2* was upregulated in the vasculature of these mature, differentiated cells now threatened by the pathogen (Fig. S3A, B). Therefore, PDF1.2 possibly only acts in mature, differentiated cells, while JA probably acts via a different pathway, such as VEGETATIVE STORAGE PROTEIN 2 (VSP2), in the undifferentiated tissue of the root tip. We next imaged *ERF1* and, like *AOS*, found it to be robustly upregulated in the same set of vascular cells that showed the *AOS* expression maximum (Fig. 4E, F). However, in contrast to *AOS*, the *ERF1* expression pattern returned to normal immediately behind the distinct group of cells that showed upregulation of *AOS*, i.e., it was moderately expressed in all cell types, not just the vasculature (Fig. 4E, F). The marker for ET biosynthesis, *ACS2*, was also upregulated in a distinct group of cells next to the colonized tissue, with its maximum partially overlapping that of *AOS* and *ERF1* (Fig. 4A-D). However, based on fluorescence intensity, *ACS2* expression appeared to be induced much stronger than *AOS* or *ERF1* (Fig. 4A-D). Further, *ACS2* expression covered both, the SA-induced HR- and the JA-induced defense response zone, with an apparent expression maximum at the mid to end of the HR zone and the boundary between the two zones. Behind the cells of the defense response zone with the expression maximum of the JA-markers, *ACS2* expression immediately dropped back to basal levels, thereby more closely resembling the pattern of *ERF1* not *AOS*. Expression remained restricted to the central cylinder for a long stretch, however, and only expanded to all cells and tissues again in the older parts of the root, close to the root’s base and the root-to-shoot transition zone. Thus, while SA and JA appear to be strictly separated spatially, ET may bridge the two zones.

**Figure 3:**
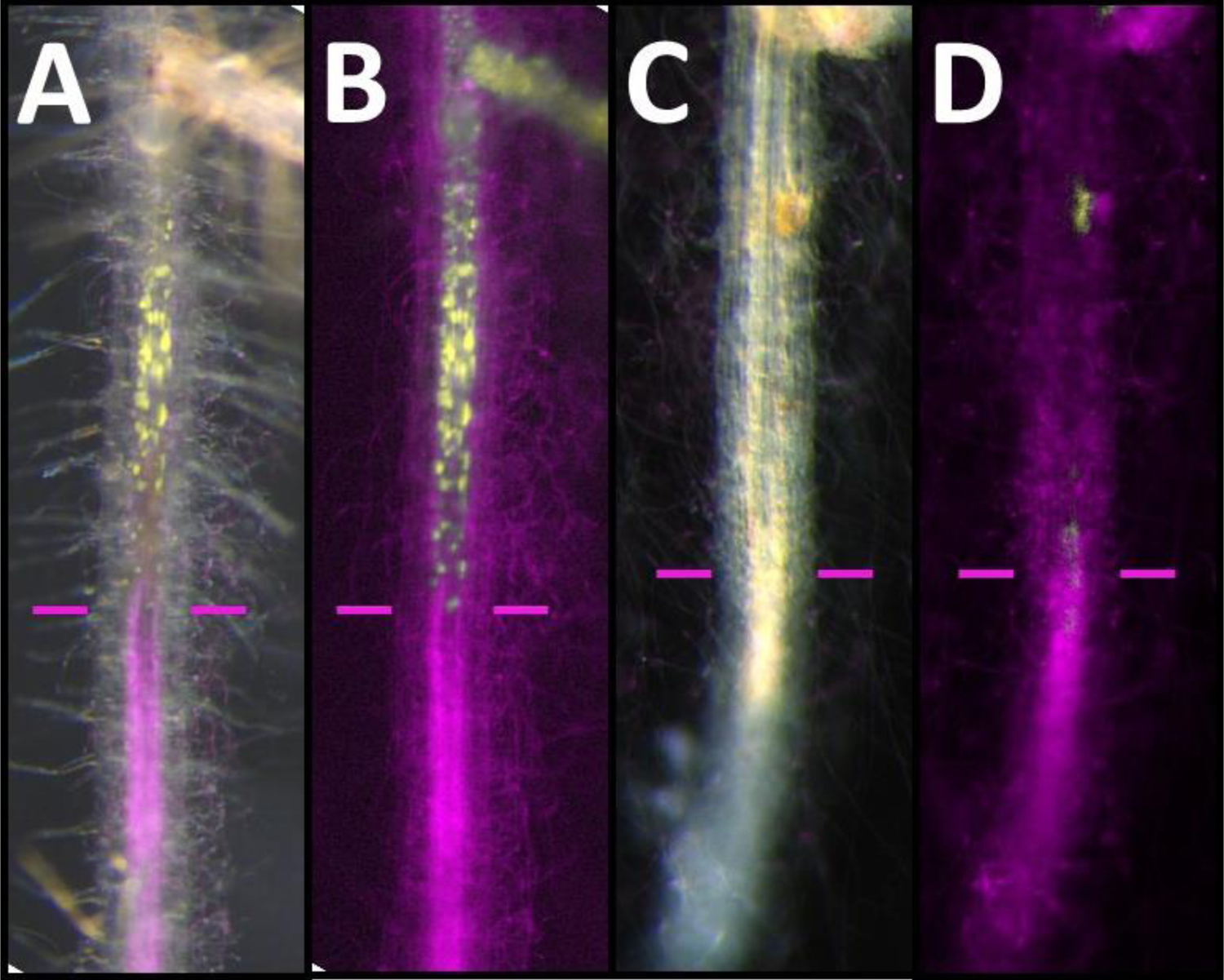
Expression of *AOS* and *PDF1.2* following colonization of the root by *Fo*5176. **A, B**: Overlay of bright field and fluorescence (A), and fluorescence-only (B) images of a root tip expressing the *AOS* reporter (yellow) in response to colonization by *Fo*5176 (magenta). **C, D**: Overlay of bright field and fluorescence (C), and fluorescence-only (D) images of a root tip expressing the *PDF1.2* reporter (yellow) in response to colonization by *Fo*5176 (magenta). The colonization front is marked by magenta bars.

**Figure 4:**
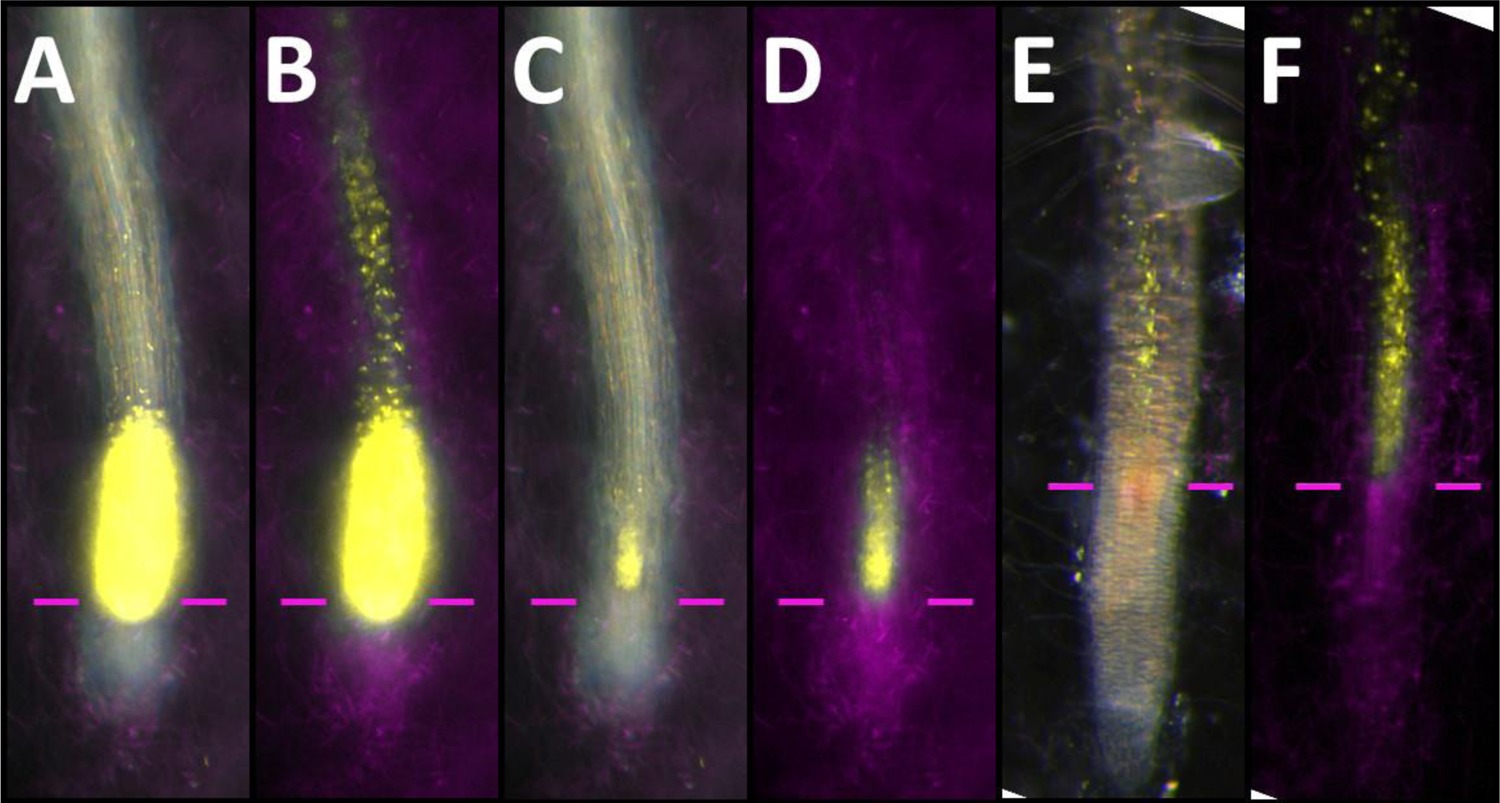
Expression of *ACS2* and *ERF1* following colonization of the root by *Fo*5176. **A, B**: Overlay of bright field and fluorescence (A, C), and fluorescence-only (B, D) images of a root tip expressing the *ACS2* reporter (yellow) in response to colonization by *Fo*5176 (magenta). The images in C and D are the same as A and B with reduced exposure time to allow for the identification of individual cells. **E, F**: Overlay of bright field and fluorescence (E), and fluorescence-only (F) images of a root tip expressing the *ERF1* reporter (yellow) in response to colonization by *Fo*5176 (magenta). The colonization front is marked by magenta bars.

### A WRKY11/WRKY70 module could coordinate the spatially distinct JA and SA responses

WRKY11 and WRKY70 act upstream of JA and SA signaling respectively, and act as integrators of these two mutually antagonistic pathways (Journot-Catalino *et al*., 2006). Therefore, the action of these two transcription factors may also establish the two distinct zones of JA and SA activity in the colonized root tip. WRKY11 is considered a negative regulator of basal plant resistance, but a positive regulator of JA biosynthesis. WRKY70, on the other hand, is a positive regulator of SA, but a negative regulator of JA. Furthermore, *WRKY70* itself is under negative regulation by WRKY11 and JA-signaling (Journot-Catalino *et al*., 2006). To test if this regulatory module is also active in response to *Fo*5176 infection, we imaged our *WRKY11* and *WRKY70* reporter lines.

We found that under control, non-infected conditions *WRKY11* is expressed at a moderate level throughout the plant in all tissues, as expected for a gene involved in negatively regulating basal resistance (Fig. 5A, B). However, upon colonization of the vasculature by *Fo*5176, the pattern of *WRKY11* expression changed, now confined to the vasculature only, with a clear expression maximum in the cells neighboring the colonized cells, overlapping the maxima of *AOS* and *ERF1* (Fig. 5C, D). Expression was removed from the surrounding tissues (epidermis, cortex), possibly to allow for the expression of basal resistance genes normally repressed by WRKY11. This enhanced expression pattern persisted for a long stretch up the root, before it reverted to normal levels in cells closer to the root base, far away from the colonization site. This long tail of expression mirrored that of *AOS* expression but was differed from that of *ACS2* or *EDS16*. Thus, transcriptional activation of JA biosynthesis and signaling indeed overlaps with the activation of its regulator WRKY11.

**Figure 5:**
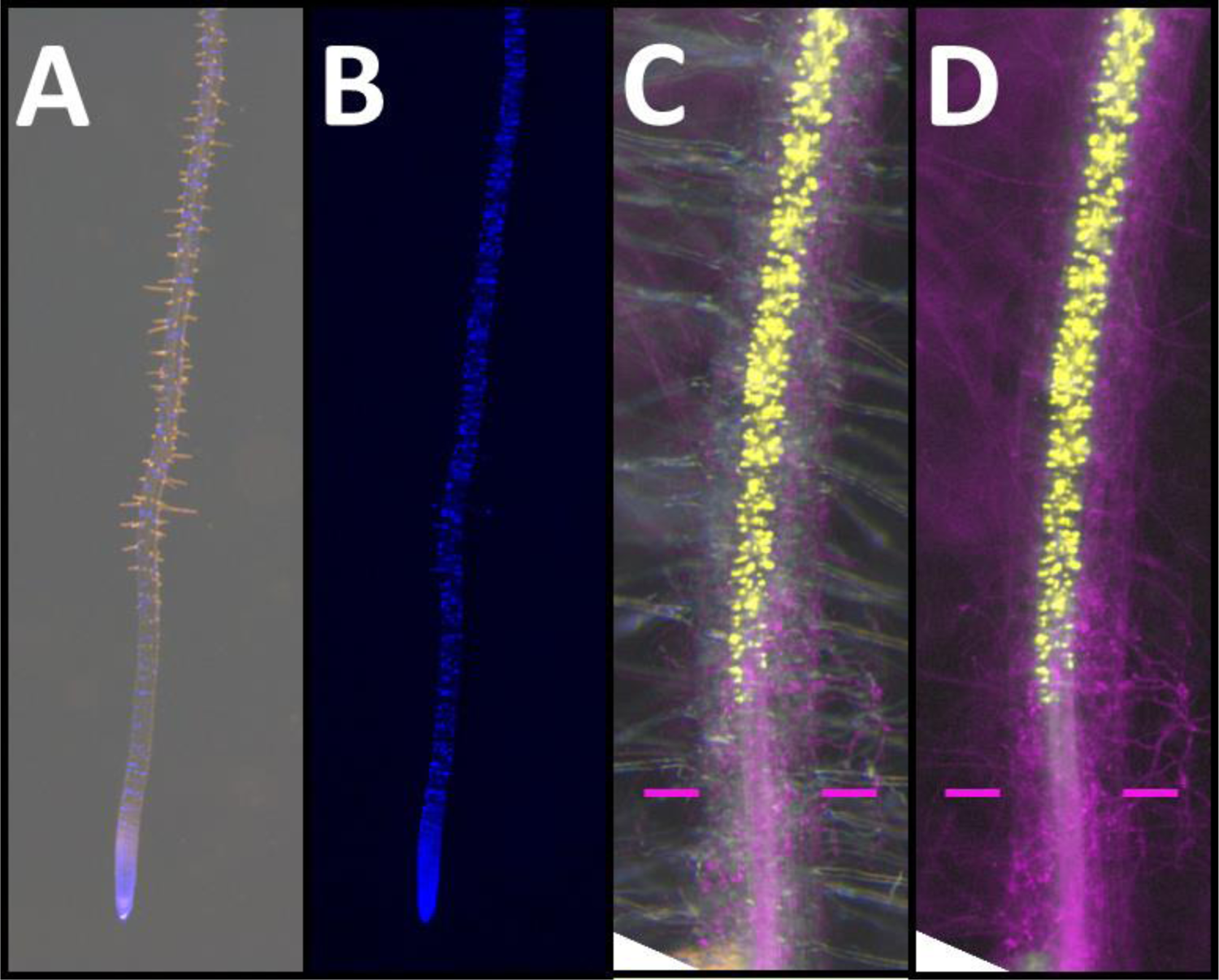
Expression of *WRKY11* in an uninfected control and following colonization of the root by *Fo*5176. **A, B**: Overlay of bright field and fluorescence (A), and fluorescence-only (B) images of an uninfected control root expressing the *WRKY11* reporter (blue). These overview images were taken with 12x magnification. **C, D**: Overlay of bright field and fluorescence (C), and fluorescence-only (D) images of a root tip expressing the *WRKY11* reporter (yellow) in response to colonization by *Fo*5176 (magenta). The colonization front is marked by magenta bars.

We then imaged the *WRKY70* reporter to see if its proposed role as a positive regulator of SA-, and negative regulator of JA-signaling is also reflected in its expression pattern in response to colonization by *Fo*5176. Under control conditions, *WRKY70* was expressed in all cells and tissues of the root, thought the expression appeared stronger in the outer tissues surrounding the central cylinder (Fig. 6A, B). Following colonization by *Fo*5176, we observed strong upregulation in vascular cells neighboring the colonized tissue, with a maximum in cells directly in contact with the colonized cells, overlapping the maximum seen for SA biosynthesis and signaling (Fig. 6C, D). This maximum was followed by a tail of residual *WRKY70* expression across the JA-zone, but in this zone, expression still appeared to be stronger in the cells surrounding the central cylinder, and therefore not directly overlapping *WRKY11* expression (Fig. 6C, D). Thus, WRKY70 may indeed be responsible for promoting SA biosynthesis and signaling in the HR zone, while the mutual antagonism of WRKY11 and WRKY70 could contribute to establishing these two distinct response zones.

**Figure 6:**
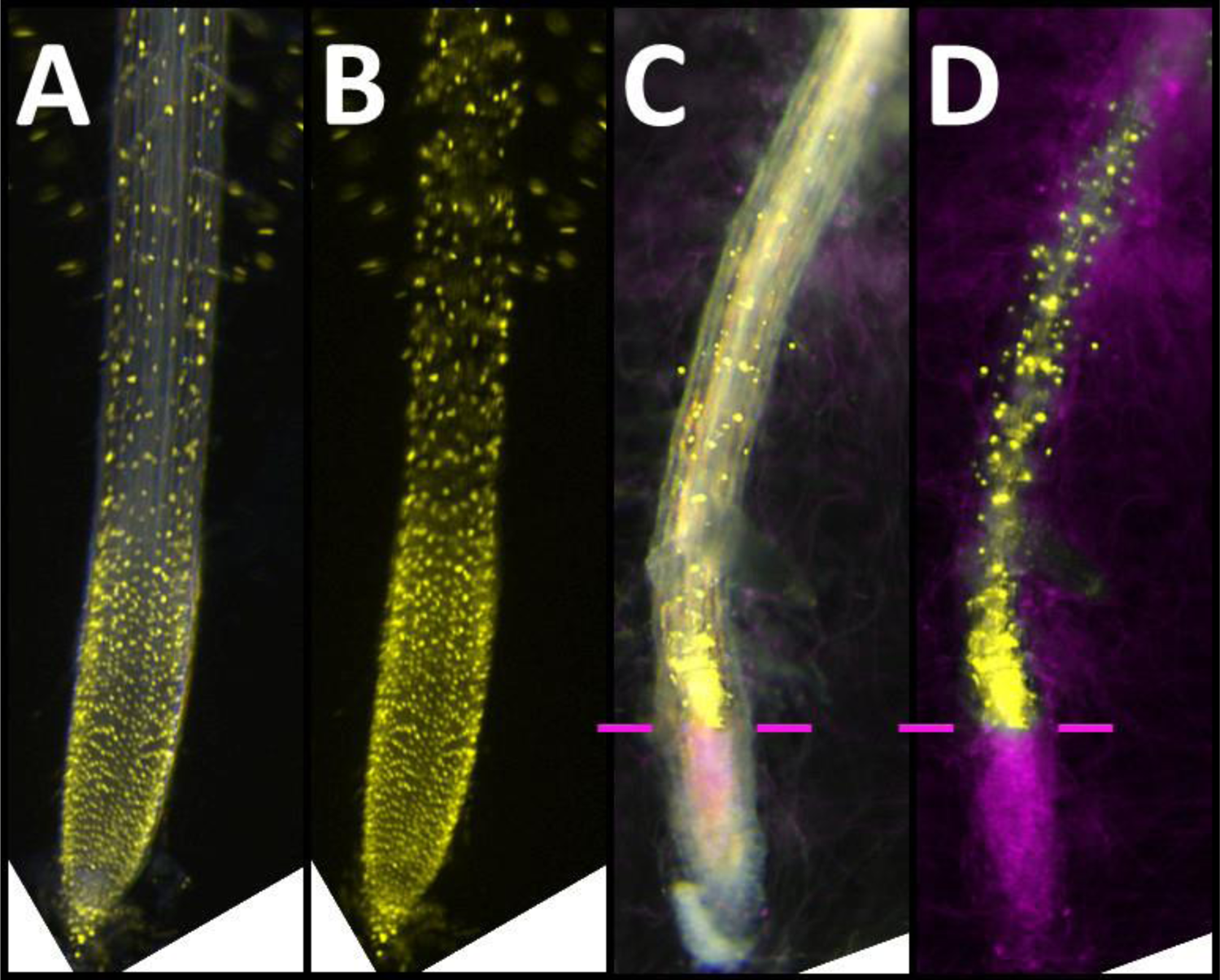
Expression of *WRKY70* in an uninfected control and following colonization of the root by *Fo*5176. **A, B**: Overlay of bright field and fluorescence (A), and fluorescence-only (B) images of an uninfected control root expressing the *WRKY70* reporter (yellow). **C, D**: Overlay of bright field and fluorescence (C), and fluorescence-only (D) images of a root tip expressing the *WRKY70* reporter (yellow) in response to colonization by *Fo*5176 (magenta). The colonization front is marked by magenta bars.

## Discussion

With the work described here we aimed at clarifying some of the inconclusive data concerning the roles of JA, ET and SA in regulating the plant’s defense against the fungal pathogen *Fo*5176. Consistent with our own microscopic observations, and previously described by Cyzmmek et al. (2007), *Fo*5176 infects and colonizes *A. thaliana* almost exclusively via the root tip. We therefore reasoned that previous bulk transcriptomic analyses using whole seedlings, or the entire root system were insufficient to detect highly specific events occurring within the limited amount of infected tissue. This hypothesis was strengthened by our initial observation, that the infected tissue undergoes immediate cell death (Fig. 1), thereby limiting the amount of infected tissue available for transcriptomic analyses even further. By employing a microscopy-based approach to live-image the plant’s response to infection in real time and at individual cell resolution, we were then indeed able to show that JA, ET and SA all contribute to the plant’s defense against *Fo*5176, albeit in clearly distinct groups of cells (Fig. 7).

**Figure 7:**
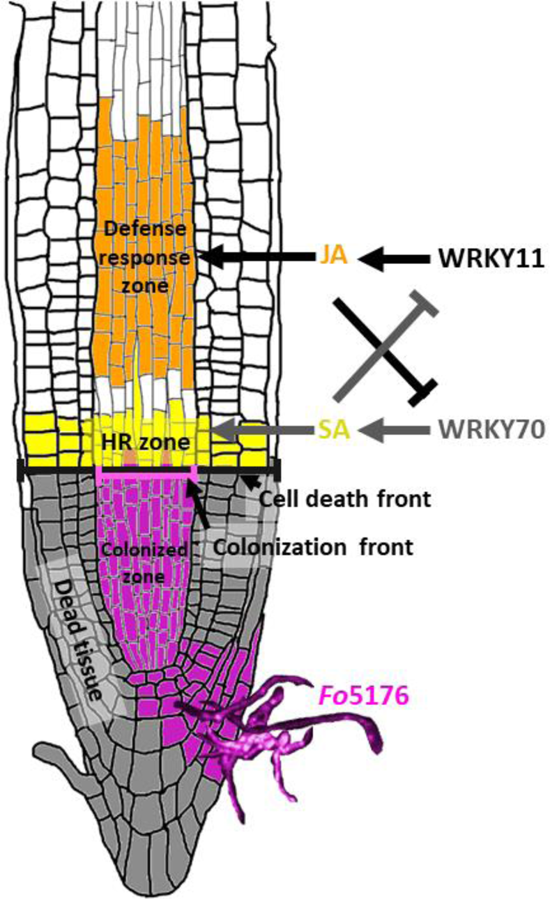
Model showing the distinct activity zones of the phytohormones and transcription factors. *Fo*5176 infects an *A. thaliana* root tip via the meristem in the tip, where it colonizes the vasculature (purple cells = colonized zone). The tissue around this zone undergoes cell death (dark grey cells). As the colonization front (purple line) progresses in the vasculature, so does a cell death front (black line) across all tissues. In an apparent attempt to prevent the fungus from infecting more tissue, the plant triggers a cell death response in a group of cells immediately adjacent to the colonization front (yellow cells), the HR zone. HR is activated by WRKY70 via SA. Slightly removed from the HR and colonized zones, the plant launches an active defense response to combat the pathogen in the defense response zone (orange cells). This response is dependent on WRKY11 and JA/ET-biosynthesis and signaling. WRKY11/JA and WRKY70/SA are mutually antagonistic, thereby establishing the two spatially separate zones of distinct action.

### An SA signaling pathway establishes the HR zone next to the colonization site

We first noticed that colonization of the vasculature was accompanied by local cell death of the entire infected root tip. As the fungus continued to move through the root, colonizing more cells, it formed a clearly visible ‘colonization front’ in the vasculature. As this colonization front moved forward, a ‘cell death front’ moved in line with the colonization front. The only difference between these two fronts being that the cell death front ran along all tissues of the root, from epidermis to epidermis, and not just in the vasculature, as is seen with the colonization front (Fig. 7). We hypothesized that this cell death front was the result of the plant’s own HR, rather than necrotrophic behavior by the fungus, and therefore imaged our reporters for SA biosynthesis and signaling, since SA is a known inducer of HR (Pitsili *et al*., 2020). Indeed, both SA biosynthesis and signaling, were upregulated precisely in a small group of cells directly in contact with the already colonized and dead tissue. Our interpretation of this observation is that SA induces HR in the cells that will otherwise be colonized next by the pathogen, thereby trying to prevent the fungus from spreading further. In our model we designated this group of cells the ‘HR-zone’ (Fig. 7). JA, on the other hand, was upregulated in a distinct group of vascular cells beginning slightly (4-7 cells) removed from the HR-zone. In this region, that we named the ‘defense response zone’, we think the plant launches an active defense response to combat the invader (Fig. 7). Such defense can include the activation of NADPH oxidases to produce ROS in the apoplast that may damage the fungus, an acidification of the apoplast via proton pumps, or the production of antimicrobial compounds (Wang *et al*., 2022a). ET activity bridges the boundary between these two zones, may facilitate boundary formation between the two zones, and also contributes to the defense response zone together with JA.

### WRKY70 acts via SA to establish the HR zone, and WRKY11 acts via JA to establish a defense response zone’

We then asked how this distinct spatial separation of the two hormonal responses can be achieved and investigated the role of a WRKY transcription factor module, consisting of WRKY11 and WRKY70 that have previously been described as controlling JA and SA signaling respectively (Journot-Catalino *et al*., 2006). In this regard, we not only found that *WRKY11*, a positive regulator of JA, was indeed upregulated in the defense response zone, while *WRKY70*, a positive regulator of SA, was upregulated in HR-zone, but we also observed that these two transcription factors altered their expression patterns. In the uninfected state, *WRKY11* was expressed in all cells and tissues of the root, in accordance with a proposed role as a negative regulator of basal defense gene expression (Journot-Catalino *et al*., 2006). This is consistent with the hypothesis that under regular growth conditions, defense genes need to be downregulated/repressed, to allow for normal development of the plant, as part of the defense-development trade-off. Following colonization of the plant by *Fo*5176, *WRKY11* was robustly upregulated in the defense response zone, likely activating JA biosynthesis and signaling. In contrast, *WRKY11* expression was no longer observed in all other tissues. This would enable the release of basal defense genes from repression and prime such tissue for a possible pathogen attack. Furthermore, WRKY11 would also contribute to the spatial separation of the defense response and HR zones, since WRKY11 and JA jointly repress SA via WRKY70. Consequently, *WRKY70* expression is excluded from the defense response zone following colonization by *Fo*5176 and most likely excludes WRKY11 and JA from the HR zone. Thus, the mutual exclusion of WRKY11 and WRKY70 appears to establish the separate defense response and HR zones (Fig. 7).

### ET-signaling and RBOHD-derived ROS may establish the boundary between HR and defense response zone

Next to WRKY11 and WRKY70, ethylene and ROS-mediated signaling may also contribute to the formation of a boundary between the HR- and defense response zones. The induction of HR results from the co-incidence of PAMP-detection via PRRs and local cell damage caused by a pathogen, which induces SA and ET biosynthesis and signaling, as well as ROS-production by RBOHD (Torres *et al*., 2005; Pogány *et al*., 2009; Lukan and Coll, 2022). Local high levels of all three factors, SA, ET and ROS, results in local cell death. Since cell death is itself a form of cell damage, this could result in a positive feedback-loop, where cell damage leads to HR, which is more cell damage leading to the further spread of HR and so on. This phenomenon is called ‘runaway cell death’ (RCD). Torres et al. (2005) have previously put forward a model to explain how RCD is prevented through the interplay of SA, ET and ROS signaling, which Pogány et al. (2009) have then expanded on (Torres *et al*., 2005; Pogány *et al*., 2009; Lukan and Coll, 2022). Sensing of a pathogen results in the upregulation of SA, ET and ROS. All three are then involved in priming the plant for impeding attack, but ROS also acts to repress SA/ET-induced HR. However, ROS can only block the SA/ET->HR pathway as long as SA and ET concentrations are at moderate levels. Acute pathogenic attack that results in cell damage elevates local SA and ET concentrations to a point at which ROS can no longer counteract it (Fig. 8). In this situation of high local concentrations of SA, ET and ROS, this co-incidence results in HR. In the case of vascular colonization by *Fo*5176, this situation would affect the cells of the HR-zone that are in immediate contact with the already colonized, and thus damaged cells. Slightly removed from the colonization front, in the cells that we call the boundary between HR and defense response zone, local SA and ET concentrations are lower, so ROS can still block the induction of HR to prevent RCD (Fig. 8). Local signaling by residual SA, JA, ET and ROS also primes these cells for impeding attack. Finally, in the defense response zone, SA is excluded, and now JA and ET jointly induce the active defense response of the plant, with ROS contributing to this defense system (Fig. 8). This model is purely hypothetical at this point, but in our previous work we could show that a ROS-generating pathway, consisting of the DAMP-acting PLANT ELICITOR PEPTIDE 1 (PEP1), its receptor PEPReceptor 2 (PEPR2), the downstream cytoplasmic BOTRYTIS-INDUCED KINASE 1 (BIK1), as well as the NADPH oxidase RBOHD, is indeed locally transcriptionally activated in response to colonization by *Fo*5176 (Calabria *et al*., 2022). Fitting with the above hypothesized role, these factors are activated in a long stretch covering both, the HR- and defense response zone, with an apparent expression maximum in the HR-zone cells and the boundary, therefore displaying a pattern similar to the one for ET biosynthesis and signaling. In this pathway, PEP1 is produced in response to pathogen-induced cell damage. This peptide is bound by the PEPR2, which triggers a phosphorylation chain from PEPR2’s intracellular kinase domain to the cytoplasmic BIK1 and the transmembrane RBOHD, which eventually produces a ROS-burst (Fig. 8) (Liu *et al*., 2013; Tintor *et al*., 2013; Holmes *et al*., 2018).

**Figure 8:**
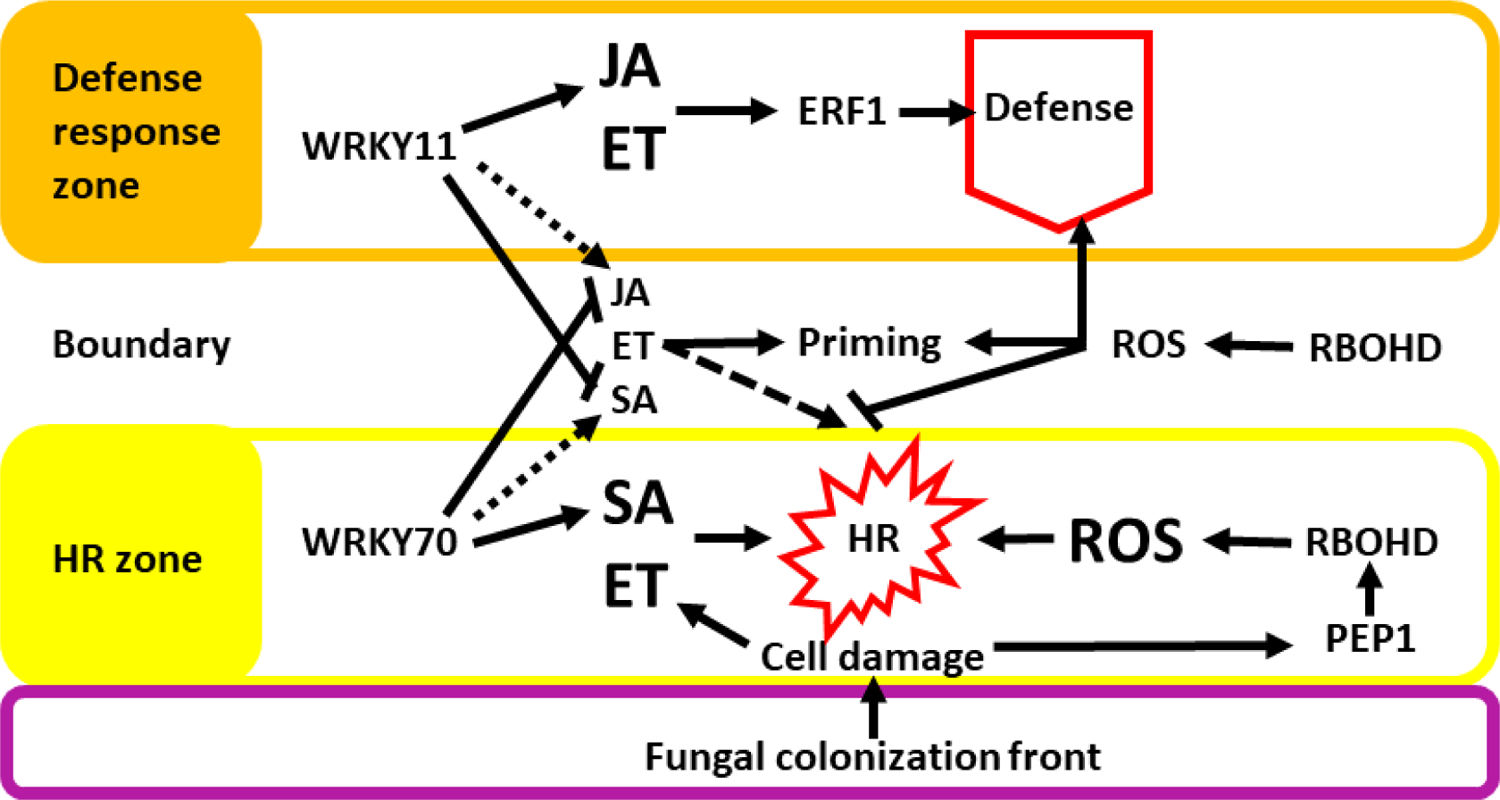
Hypothetical model of how the phytohormones and the PEP pathway may establish the different zones and boundary. In the cells of the HR zone, PEP1, via PEPR2 and BIK1, activates RBOHD to produce ROS. In combination with an over-accumulation of SA and ET via WRKY70- and cell damage-signaling, these factors induce HR. In the boundary, SA and ET accumulate at lower concentrations because the cell damage signaling is lacking and WRKY11 signaling partially suppresses WRKY70 signaling. Thus, ROS can now suppress HR and thus RCD. In the cells of the defense response zone, WRKY11 activates JA biosynthesis, while ET is still active, potentially via PEP1 signaling. JA and ET jointly activate active defense via ERF1.

### Limitations of this study and outlook

Whilst the inclusion of the WRKY-module and the dual role of RBOHD in contributing to defense and controlling RCD led to an elegant hypothesis of how the separate zones can be established, further work is needed to validate these observations. First, it will be important to image the phytohormone markers in *wrky11* and *wrky70* mutant plants, to definitively establish that they act upstream of the hormone pathways. The investigation of JA biosynthesis and signaling mutants, as well as the SA-draining *NahG* transgenic line, will furthermore inform us more closely on the supposed role of these hormones (active defense vs. HR). Furthermore, while we found that the HR-zone spreads out across all tissues of the root, we only found WRKY70 and SA upregulation in the vasculature, not in the outer tissues. Thus, it is not clear what actually induces the HR in these cells. This could, however, be a normal response to a dying vasculature. Should the vasculature undergo cell death, it is conceivable that the loss of conductivity and structural integrity of the inner cylinder of the root spontaneously induces cell death of the outer tissue, without the need of active SA-signaling. Then, *rbohd* and ethylene biosynthesis mutants need to be investigated for boundary formation, and the inclusion of RBOHF may help differentiate between ROS production involved in HR and boundary formation, and ROS involved in active defense. It will be interesting to further investigate the different pathways activated specifically in the defense response and HR zones.

It is therefore clear that the two models presented here in the discussion (Figs. 7 & 8) are still very much speculative until we have completed the mutant analysis. However, we believe that the work reported here is already an interesting and informative proof-of-principle to demonstrate how a microscopy-based approach at resolving the spatiality of the plant’s immune response can add to our understanding of the plant immune system. The fact that the highly targeted and non-overlapping responses zones for JA and SA in response to colonization by *Fo*5176 have so far been overlooked, highlights the need for further high-resolution analysis to resolve the plant’s immune response to pathogenic infection on an individual cell level within intact tissue. By building on these initial results, and in combination with the pGG-PIP promoter collection that we have published and donated to AddGene (Calabria *et al*., 2022), it should be possible to further map the distinct spatial and temporal responses of other plant pathways to infection by *F. oxysporum* and any other pathogen, thereby creating a root atlas of the plant’s immune system.

## Material & Methods

### Cloning of *A. thaliana* reporter constructs

All expression clones for the different reporter lines were assembled as in Calabria *et al*. (2022), and the putative promoter sequences are from our pGG-PIP vector collection (Calabria *et al*., 2022). The expression clones are based on the pGGZ003 GreenGate destination vector backbone from (Lampropoulos *et al*., 2013). GreenGate modules for the Nuclear Localization Signal (pGGD007), UBQ terminator (pGGE009) and hygromycin resistance cassette (pGGF005) were also from the original GreenGate kit (Lampropoulos *et al*., 2013). The B- and C-modules with *mTurquoise2* (amplified from an AddGene-derived template (Goedhart *et al*., 2012)), as well as the B-module with *mNeonGreen* (amplified from an AddGene-derived template (Shaner *et al*., 2013)) and the C-module with *LTI6b* (amplified from total cellular *A. thaliana* Columbia DNA) were created using the pGGB000, pGGC000 and pGGD000 entry clones from the original GreenGate kit (Lampropoulos *et al*., 2013). Expression clone assembly was done with the NEB Golden Gate Assembly Kit (BsaI-HF^®^ v2).

### Plant growth and transformation

*A. thaliana* natural accession Columbia (Somssich, 2018) plants were grown until flowering stage at day/night conditions of 21 °C/18 °C, 16 h/8 h with a light intensity of 120 mmol m^-2^ s^-1^ and constant relative humidity of 70%. Plants were transformed using a slightly modified *Agrobacterium tumefaciens*-mediated floral dip protocol (Clough and Bent, 1998; Somssich, 2019). *A. tumefaciens* strain GV3101 *pMP90 pSoup* carrying the plasmids of interest were grown in 250 mL of 2×YT media supplemented with 50 µg/ml rifampicin, 25 µg/ml gentamycin and 100 µg/ml spectinomycin for 16 h at 28 °C, shaking at 200 rpm (Miller, 1972; Holsters *et al*., 1980; Koncz and Schell, 1986; Hellens *et al*., 2000). *A. tumefaciens* cultures were centrifuged at 3750 rpm for 20 min, and the pellet resuspended in 300 mL of 5% sucrose solution with 0.08% Silwet L-77 (Clough and Bent, 1998). Plants were inverted and the flowers submerged in *A. tumefaciens* solution for 30 s with agitation. Plants were briefly patted dry to remove excess *A. tumefaciens* before laying horizontal in a plant tray covered with plastic film to maintain humidity at room temperature overnight. The following day, plants were reverted upright and returned to the original growth conditions. The floral dipping procedure was repeated after a 7-day interval and upon maturation of siliques, seeds were harvested. Harvested seeds were dried for at least two weeks and then surface-sterilized by washing them for 2-3 h in 75% ethanol with 0.1% Triton X-100 on an upright tube rotator. The seeds were then decanted onto filter paper in a laminar flow cabinet to evaporate the ethanol. The dry seeds were then sprinkled onto a petri dish with half-strength basal Murashige & Skoog (MS) medium with vitamins supplemented with 30 µg/ml hygromycin (Murashige and Skoog, 1962). Following a 3–5-day stratification period in darkness at 4 °C, the plates were placed into a growth cabinet and grown for 10 to 14 days. After this, seedlings that survived selection were transferred to fresh plates without antibiotics and grown until they were too big for the plates, at which point they were transferred to soil until maturation of siliques.

### Fungal growth and transformation

The *Fo*5176 line expressing the cytoplasmic tdTomato (tdT) was described in our previous paper, Calabria *et al*. (2022). It was created using plasmid pMAI32, carrying a constitutive promoter from the *actin* gene from ascomycete *Leptosphaeria maculans* (Idnurm *et al*., 2017). The *tdT* coding sequence was amplified from an AddGene-derived template and inserted into pMAI32 via Gibson assembly (Shaner *et al*., 2004). The resulting expression clone was used to transform *A. tumefaciens* strain EHA105, and positive colonies were selected on LA plates supplemented with 50 µg/ml kanamycin. *Fo*5176 was cultured on potato dextrose agar (PDA) plates (Idnurm *et al*., 2017). To obtain spores for transformation, a 50 ml half strength potato dextrose broth was inoculated with five 5 mm plugs from the PDA plates. This culture was incubated at room temperature while shaking at 150 rpm for five days, then filtering through miracloth, and the spores harvested via centrifugation at 3000 rcf for 5 min. The spores were then resuspended in sterile water. For the transformation, the *A. tumefaciens* strain was cultured overnight in LB supplemented with kanamycin, and fungal spores and bacteria were then mixed on induction medium plates and cocultured for three days (Idnurm *et al*., 2017). For selection of positive transformants, an overlay of PDA supplemented with 50 µg/ml hygromycin, and 100 µg/ml cefotaxime was used. The transformants that grew through the overlay were subcultured on PDA containing hygromycin and cefotaxime and allowed to produce conidia. The final strain was then derived from a single conidium.

### Plant-fungus co-cultivation and infection

Plant-fungus co-cultivation was done as in Calabria et al. (2022), which uses an adapted protocol from Tintor et al. (2020): *Fo*5176 was routinely maintained on PDA plates at room temperature. For each co-cultivation experiment, five 5 mm plugs were cut from the plate to inoculate a 50 ml yeast nitrogen base (YNB) medium supplemented with 1% sucrose, and this culture was incubated at room temperature for four days, shaking at 120 rpm. Spores were harvested by filtering through mirocloth and centrifugation at 3000 rcf for 10 min and were then resuspended in 25 ml autoclaved MilliQ water. The *A. thaliana* plants were grown on vertical plates with half-strength basal MS medium with vitamins for 11 days, and then transferred to a horizontal plate, in which the MS medium was removed, except for a roughly 2-3 cm strip at the top (Tintor *et al*., 2020). The rest of the plate was filled up with quarter-strength liquid basal MS medium with vitamins. The seedlings were places onto the narrow solid medium strip, with the roots hanging into the liquid medium. Fungal spores were then added into the liquid medium. The plates were wrapped in aluminum foil, with only the above-ground organs exposed to light and placed back into the growth chamber.

### Imaging

We observed earliest infection of *A. thaliana* by *Fo*5176 at three days post inoculation (dpi) with spores, so we imaged seedlings and fungi daily from 3 dpi to 11 dpi. Imaging was done on a Leica M205 FA stereomicroscope using ET CFP (ET436/20x ET480/40m), ET mCHER (ET560/40x ET630/75m) and F/AL488 (ET480/40x ET535/50m) filters for the detection of mTurquoise2, tdTomato and mNeonGreen, respectively. A magnification of 80× was used unless noted otherwise in the figure legend, and imaging and detection conditions, such as illumination power, exposure time, gain, etc. were kept constant across all measurements, to make images comparable and allow for at least a semi-quantitative assessment of expression strength of the different reporters. Images were taken with the Leica Application Suite software, then exported and processed using Fiji and the GNU image manipulation program (GIMP) (Schindelin *et al*., 2012).

## Acknowledgements

This work was funded by the Australian Research Council (grant no. DE200101560) and was further supported by a seed grant from the Melbourne University Botany Foundation. LW was supported by the China Scholarship Council. H-WC was supported by a Graduate Research Scholarship from the University of Melbourne. Imaging was done on instruments maintained by the Biological Optical Microscopy Platform (BOMP) and the BioSciences Microscopy Unit at the University of Melbourne. We would like to thank Dr. Imre E. Somssich for critical reading of the manuscript and helpful comments and suggestions.

## Supplementary Figures

**Figure S1:**
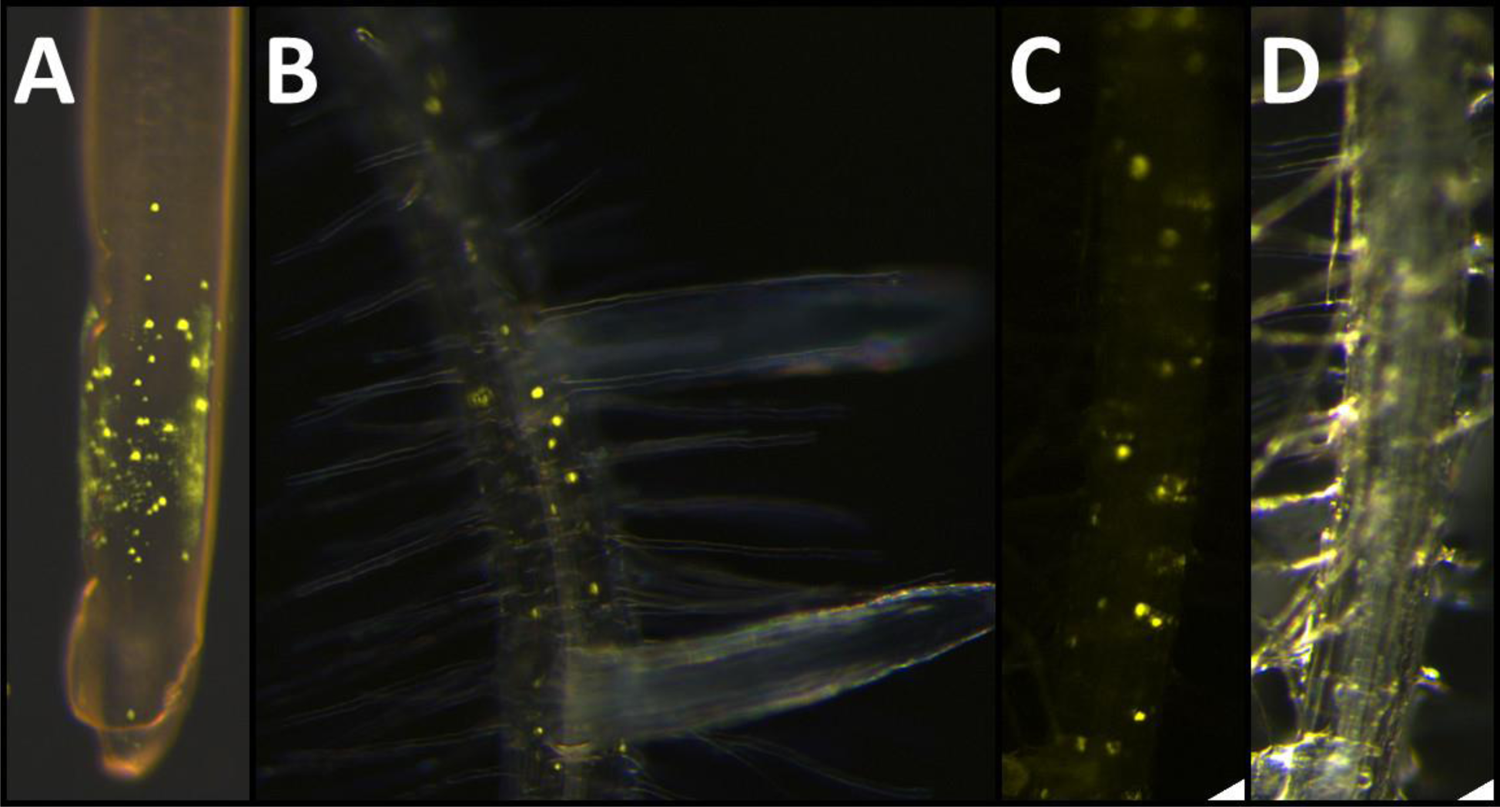
Expression patterns of *EDS16* and *PR1* in uninfected control plants. **A, B**: Overlays of bright field and fluorescence images showing fluorescence of the *EDS16* reporter in a root tip (A) and in differentiated cells (B). **C, D**: Fluorescence (C) and bright field (D) image showing fluorescence of the *PR1* reporter in differentiated cells.

**Figure S2:**
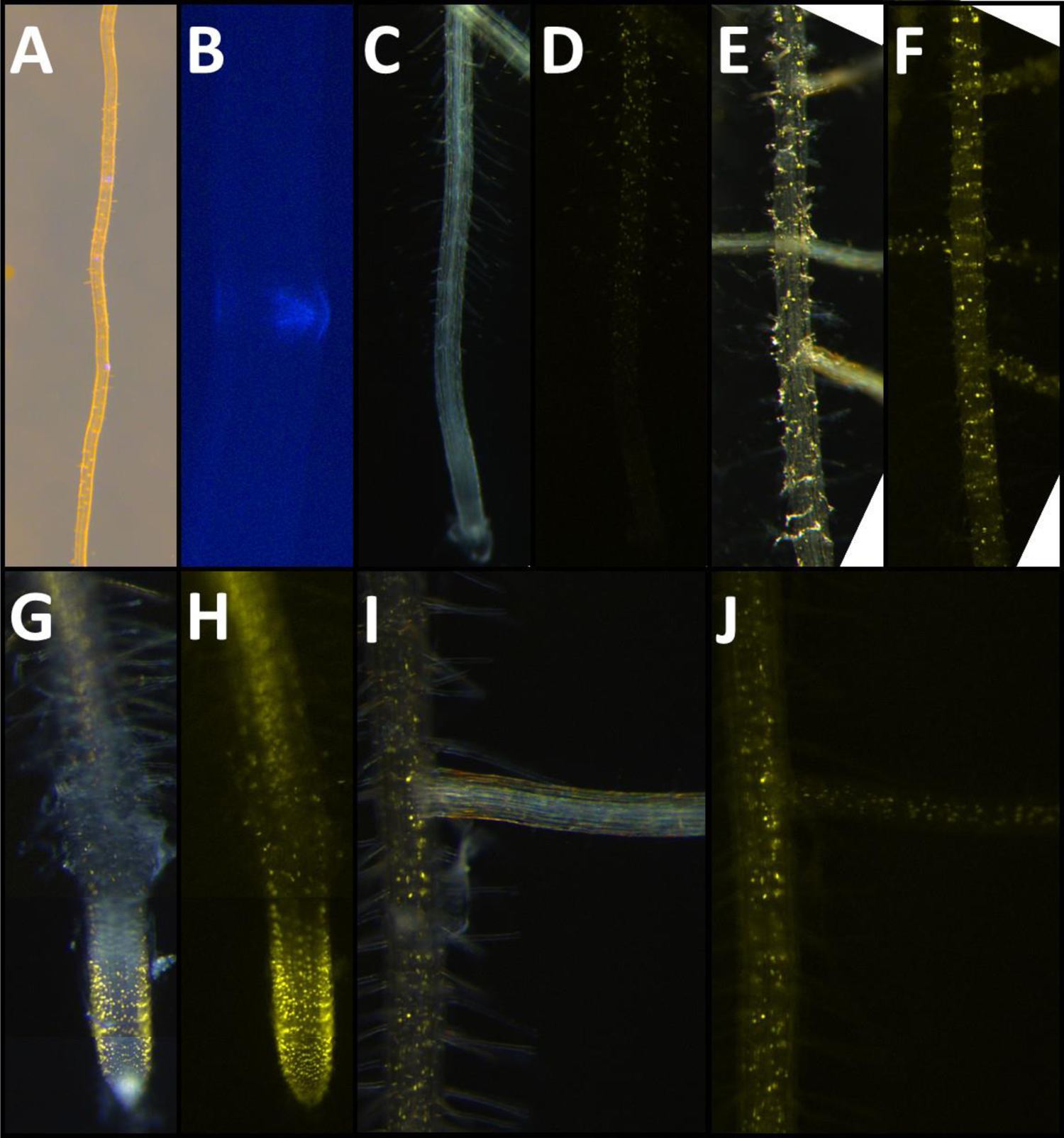
Expression patterns of *AOS*, *ERF1,* and *ACS2* in uninfected control plants. **A**: Overlay of bright field and fluorescence image showing expression of *AOS* at lateral root emergence sites. 12x magnification. **B:** Magnification of such a site. **C, D**: Overlay of bright field and fluorescence (C) and fluorescence-only (D) images showing onset of *ERF1* expression at the start of the differentiation zone. 32x magnification. **E, F**: Overlay of bright field and fluorescence (E) and fluorescence-only (F) images showing fluorescence from the *ERF1* reporter in differentiated cells. **G, H**: Overlay of bright field and fluorescence (G) and fluorescence-only (H) images showing expression of *ACS2* in the root tip and young differentiating cells. Note that these images are composites of three different images to account for a changing focal plane. **I, J**: Overlay of bright field and fluorescence (I) and fluorescence-only (J) images showing fluorescence from the *ACS2* reporter in differentiated cells.

**Figure S3:**
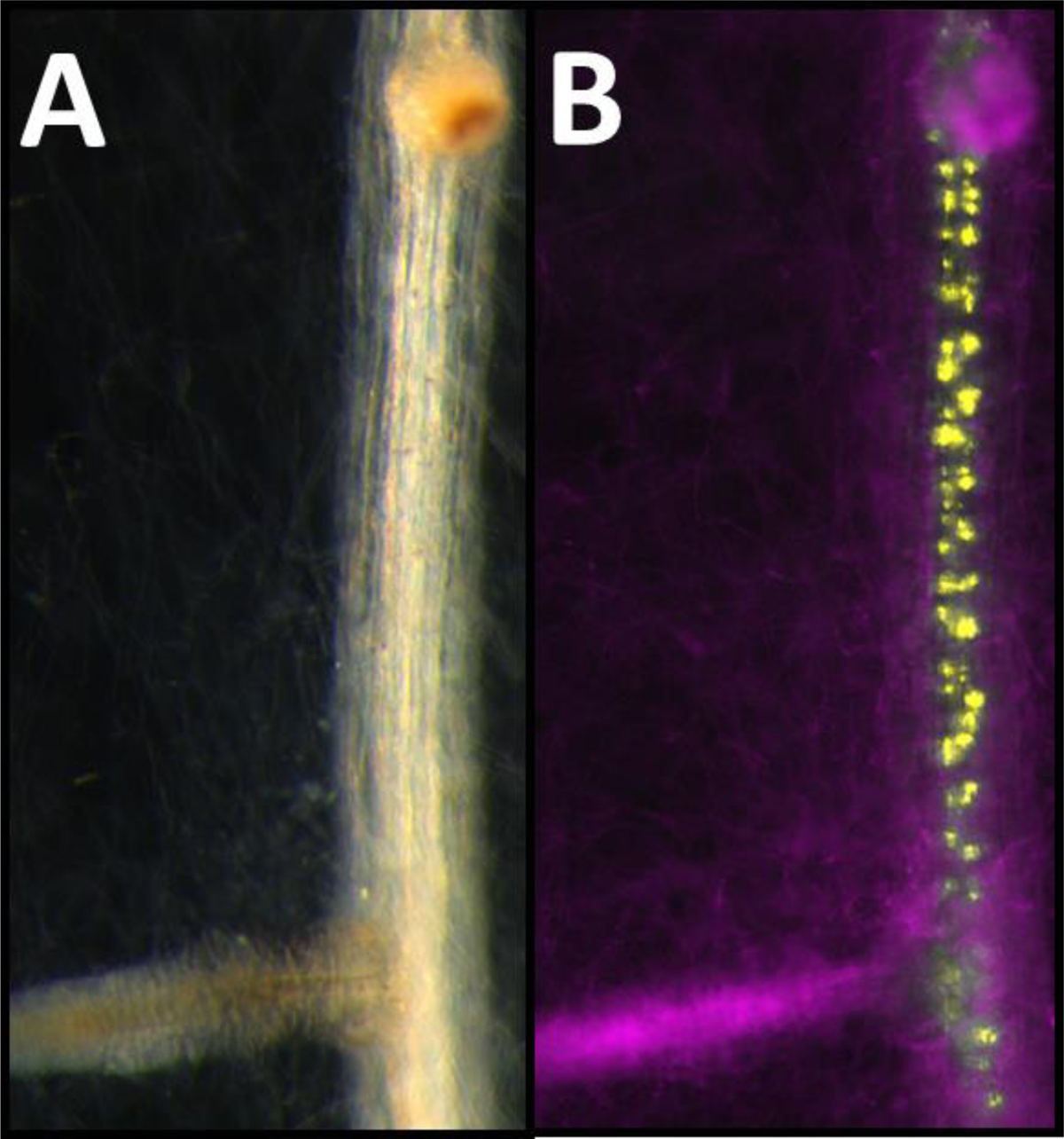
Expression of *PDF1.2* in differentiated tissue following colonization of the root by *Fo*5176. **A, B**: Bright field (A), and fluorescence (B) images of the differentiation zone of a root expressing the *PDF1.2* reporter (yellow) in response to colonization by *Fo*5176 (magenta). Colonization via the tips of lateral roots has reached the main root at its differentiation zone. In this zone *PDF1.2* is responsive to the infection, showing upregulation in the vasculature.

